# Interpersonal brain synchronization during face-to-face economic exchange between acquainted dyads

**DOI:** 10.1101/2021.12.20.473563

**Authors:** Yuto Kikuchi, Kensuke Tanioka, Tomoyuki Hiroyasu, Satoru Hiwa

## Abstract

Interpersonal brain synchronization (IBS) has been observed during social interactions and involves various factors, such as familiarity with the partner and type of social activity. A previous study has shown that face-to-face interactions in pairs of strangers increase IBS. However, it is unclear whether this can be observed when the nature of the interacting partners is different. Herein, we aimed to extend these findings to pairs of acquaintances. Neural activity in the frontal and temporal regions was recorded using functional near-infrared spectroscopy hyperscanning. Participants played an ultimatum game that required virtual economic exchange in two experimental settings: the face-to-face and face-blocked conditions. Random pair analysis confirmed whether IBS was induced by social interaction. Contrary to the aforementioned study, our results did not show any cooperative behavior or task-induced IBS increase. Conversely, the random pair analysis results revealed that the pair-specific IBS was significant only in the task condition at the left and right superior frontal, middle frontal, orbital superior frontal, right superior temporal, precentral, and postcentral gyri. Our results revealed that face-to-face interaction in acquainted pairs did not increase IBS and supported the idea that IBS is affected by “with whom we interact and how.”

## 1. Introduction

Face-to-face communication plays a vital role in building trust and cooperation with others. Cooperative behavior is also found in other animals, but large-scale and stable cooperative behavior is unique to humans (Boyd et al., 2009). What makes such a cooperation possible is the unique human ability to empathize with others and to infer their feelings and intentions (Declerck et al., 2018). It is essential to investigate how our brain functions are associated with social interaction to better understand human sociality.

In the recent years, hyperscanning, a method of simultaneously measuring the brain activity of two or more people, has been used in many studies to investigate the neural basis of social interaction (Dumas et al., 2011). It has been successfully used in various neuroimaging modalities, such as functional magnetic resonance imaging (fMRI), functional near-infrared spectroscopy (fNIRS), and electroencephalography (Salazar et al., 2021; Koike et al., 2019; Osaka et al., 2015; Chabin et al., 2020; Ciaramidaro et al., 2018). Many researchers have used these modalities to investigate various social behaviors, such as cooperation, competition, empathy, and trust. Various experimental paradigms have been proposed to experimentally reproduce such social behavior, including coordination tasks in which participants synchronize their movements (Hirsch et al., 2017; Pan et al., 2017), eye contact and joint attention tasks (Saito et al., 2010; Bilek et al., 2015), economic game tasks such as the prisoner’s dilemma and ultimatum games (Zhang et al., 2020; Tang et al., 2019), and tasks involving everyday activities such as choral singing and discussions (Osaka et al., 2015; Müller et al., 2019). Even though the experimental paradigm is different, previous studies have confirmed that interpersonal brain synchronization (IBS) in the dorsolateral prefrontal cortex and the temporoparietal junction (TPJ) is the key feature of social behavior.

Gvirts et al. (2020) proposed that IBS during social interaction can be affected by three factors: the type of social activity, the setting of the interaction, and the nature of the interacting partner. Among these factors, Tang et al. (2016) focused on the setting of interactions, investigating its effect on IBS using an ultimatum game under two different environmental settings: the face-to-face (FF) and face-blocked (FB) conditions. They found that FF interactions increased “shared intentionality,” a positive belief in each other’s cooperative decision-making, compared with FB interactions, and facilitated cooperation between partners. In addition, fNIRS-based hyperscanning demonstrated that IBS in the right TPJ was greater in FF interactions than in FB interactions and was also increased by the existence of shared intentionality between partners. Their work is significant, as they demonstrated IBS between two people engaged in FF interactions. However, it should be noted that these findings were confirmed for pairs of strangers, and it remains unclear whether these results can be reproduced when the nature of the interacting partners is different. It is also of great interest whether the nature of the interacting partners affects the increased IBS in FF interactions.

Herein, we investigated how FF interaction affects IBS in pairs of acquaintances. We used the same experimental paradigm as Tang et al. (2016), intending to extend their findings in stranger pairs to acquainted interacting partners. We investigated the following hypotheses: (1) FF interaction enhances the shared intentionality between pairs and promotes cooperation because it provides greater visual information about the partner than FB interaction; (2) IBS in the right TPJ, responsible for inferring the partner’s intentions, is greater in FF interaction than in FB interaction. These hypotheses are the same as in the original study, and we aimed to replicate their findings in acquainted pairs. In addition to the original study, we examined whether IBS was specific to the interacting pair. One of the pitfalls of hyperscanning is that an increased IBS might be observed when a different participant performs the same task independently, although there is no interaction between pairs. Therefore, we confirmed that IBS would be higher in interacting pairs than in non-interacting pairs by performing random-pair analysis.

## 2. Materials and Methods

### 2.1 Participants

Forty-eight healthy university students (24 same-sex pairs; FF: 12 pairs with 10 pairs of men, age [mean ± standard deviation]: 22.00 ± 1.12; FB: 12 pairs with 10 pairs of men, age: 22.46 ± 1.41) from Doshisha University participated in our study. All pairs belonged to the same laboratory and were acquainted before the experiment; they recognized each other’s names and faces and had daily conversations. All participants were informed of the methods and risks of the experiment and provided written informed consent. This study was conducted per the research ethics committee of Doshisha University, Kyoto, Japan (approval code: 21001).

### 2.2 Procedure and behavioral data acquisition

Each participant sat at a table with a keyboard and a display. Each pair was randomly assigned to one of two experimental environments: FF or FB. In the FF condition, the pair could see each other’s faces (Fig. 1A). The Proposer communicated only information about the reward amount and the offer verbally. The Responder was instructed not to say anything at all. In the FB condition, the pairs were separated by a partition wall and could not see each other’s faces. Under both conditions, the decisions in the task were performed by pressing the keyboard.

**Fig. 1.**
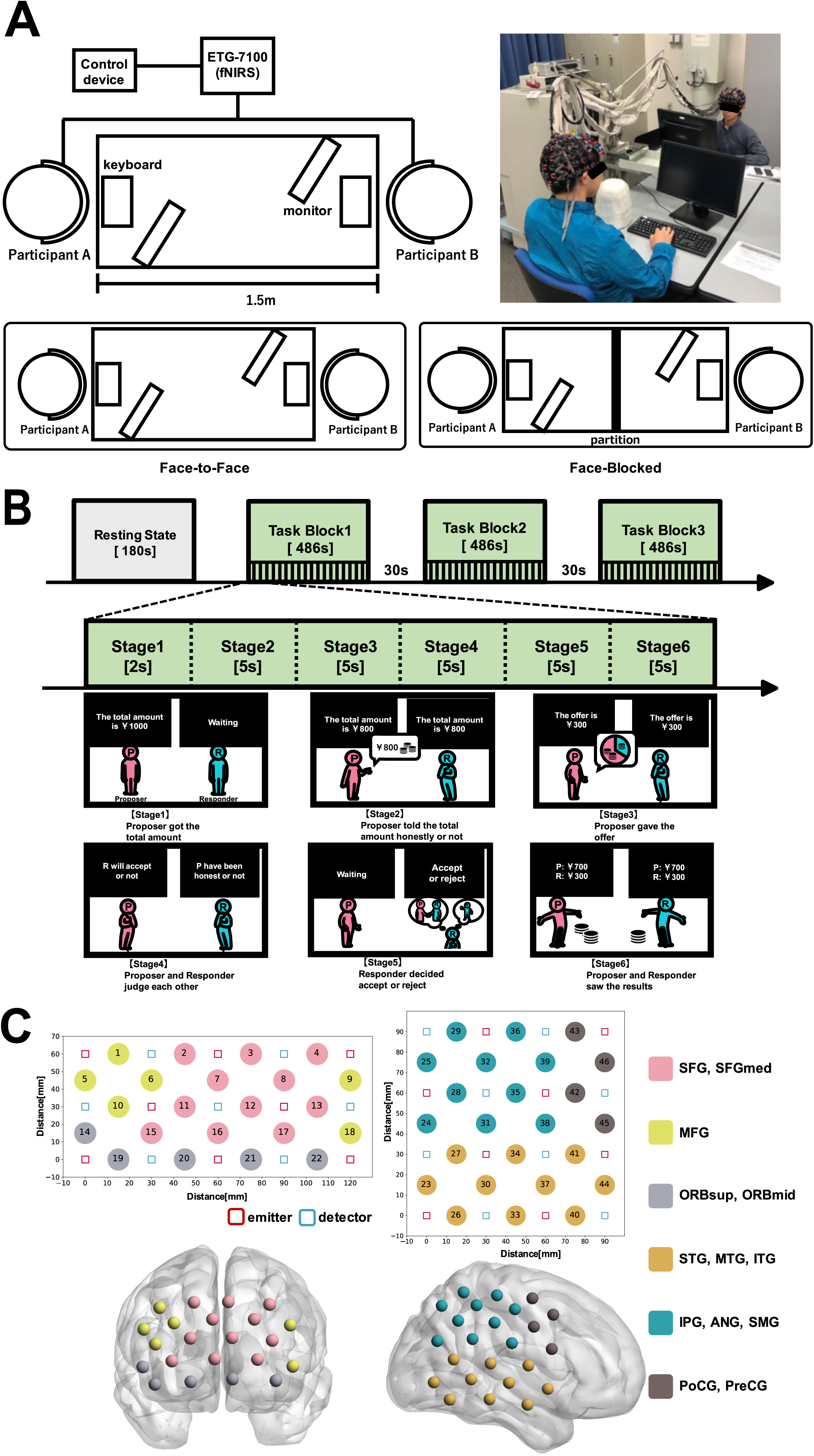
Experimental design and setup. (A) Experimental environment. FF condition: the pairs were opposite each other and could see each other’s faces. FB condition: the pairs were opposite each other but separated by a partition wall. (B) Timeline of the experimental protocol. Resting-state block: participants were instructed to be relaxed with their eyes closed and remain as still as possible. Task block: one task block consisted of 18 consecutive trials. In one trial, the pairs played one ultimatum game. There were three task blocks, and consequently, the pairs performed 54 rounds of the ultimatum game in total. (C) fNIRS probe assignment. The emitters and detectors are indicated as red and blue squares, respectively. The measurement channels were marked as circles and numbered. Anatomical regions were determined by the virtual registration method. See Table 1 for spatial registration of the fNIRS channel location to the automated anatomical labeling atlas in MNI space Abbreviations: FF: face-to-face; FB: face-blocked; fNIRS: Functional near-infrared spectroscopy; MNI: Montreal neurological institute

**Fig. 2.**
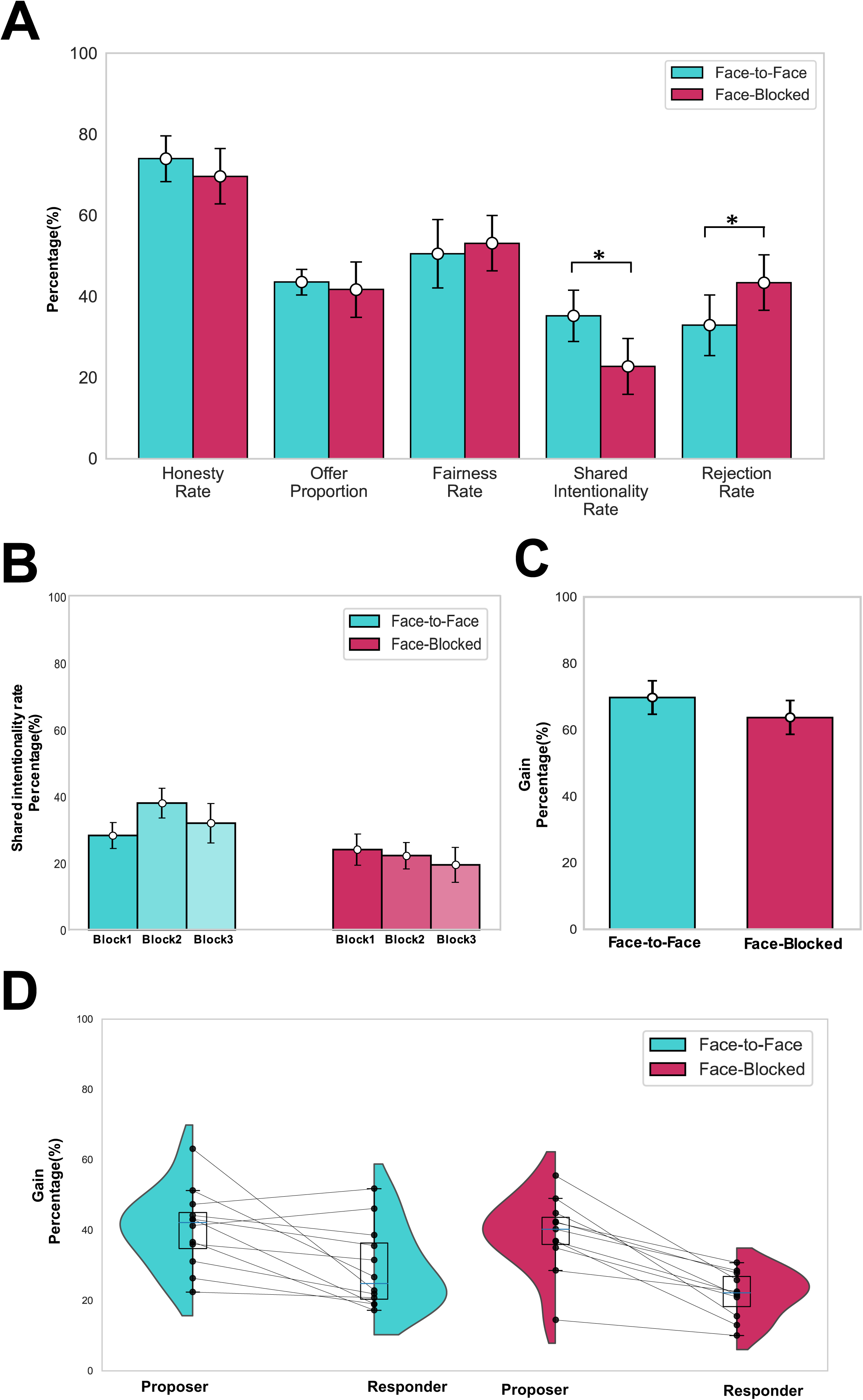
Behavioral analysis results. (A) Game measures. The SIR and RR were significantly different between the FF and FB conditions (*p* < 0.05), but the FR, OP, HR were not significantly different. Error bars indicate standard errors. (B) Time effect of SIR. There was no significant difference between the task blocks in either the FF or FB conditions. Error bars indicate standard errors. (C) Gains in the game of Proposer and Responder. There was no significant difference between the Proposer and Responder’s gains between the conditions. (D) Total amount earned for each role. In both conditions, the gains were significantly higher for the Proposer than for the Responder (FF: *p* = 0.012; FB: *p* = 2.7 × 10^-4^). Black lines connect the values of the pairs that participated in the experiment together. Abbreviations: FF: face-to-face; FB: face-blocked; FR: fairness rate; OP: offer proportion; HR: honesty rate; SIR: shared intentionality rate

**Table 1.**
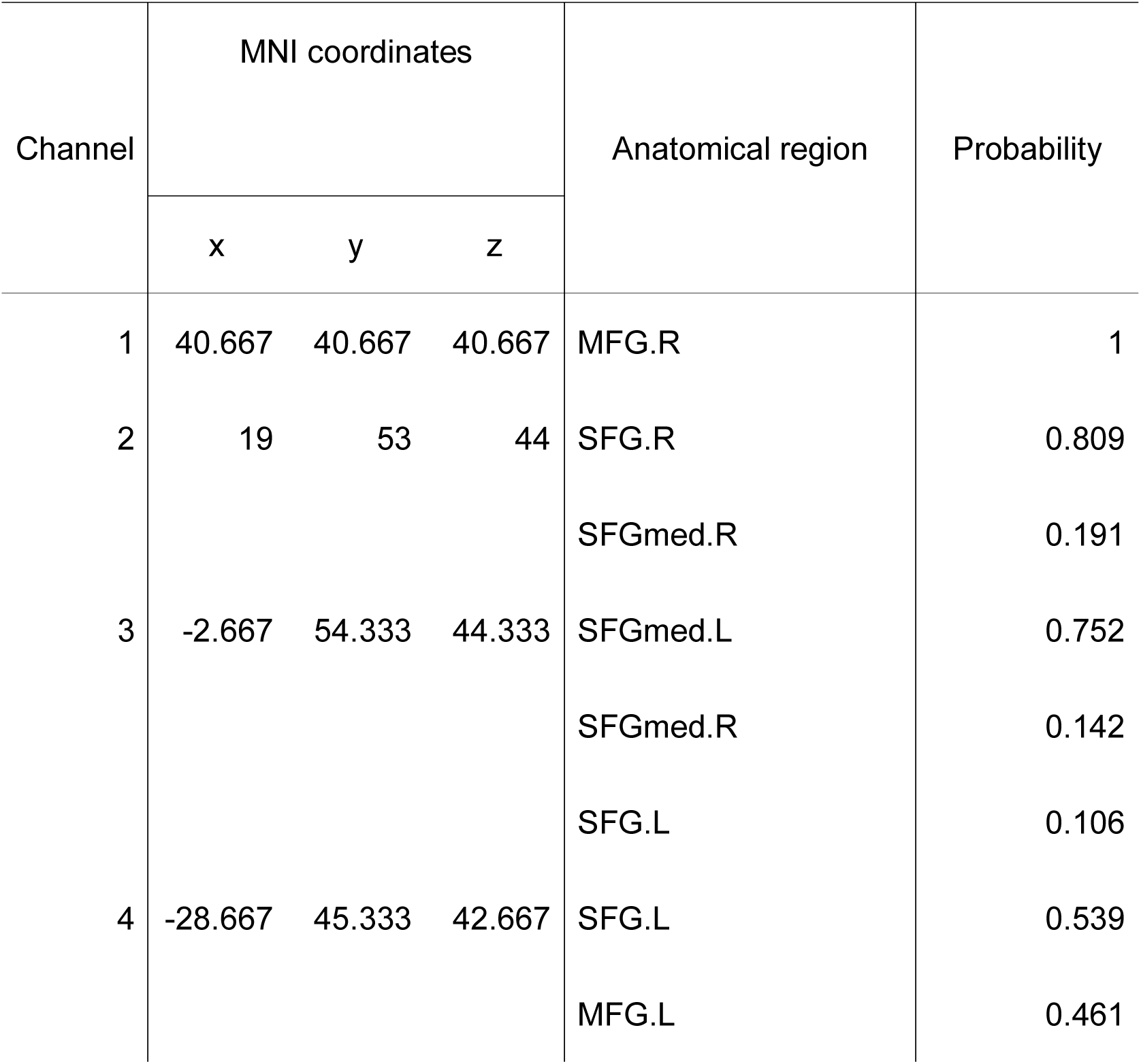
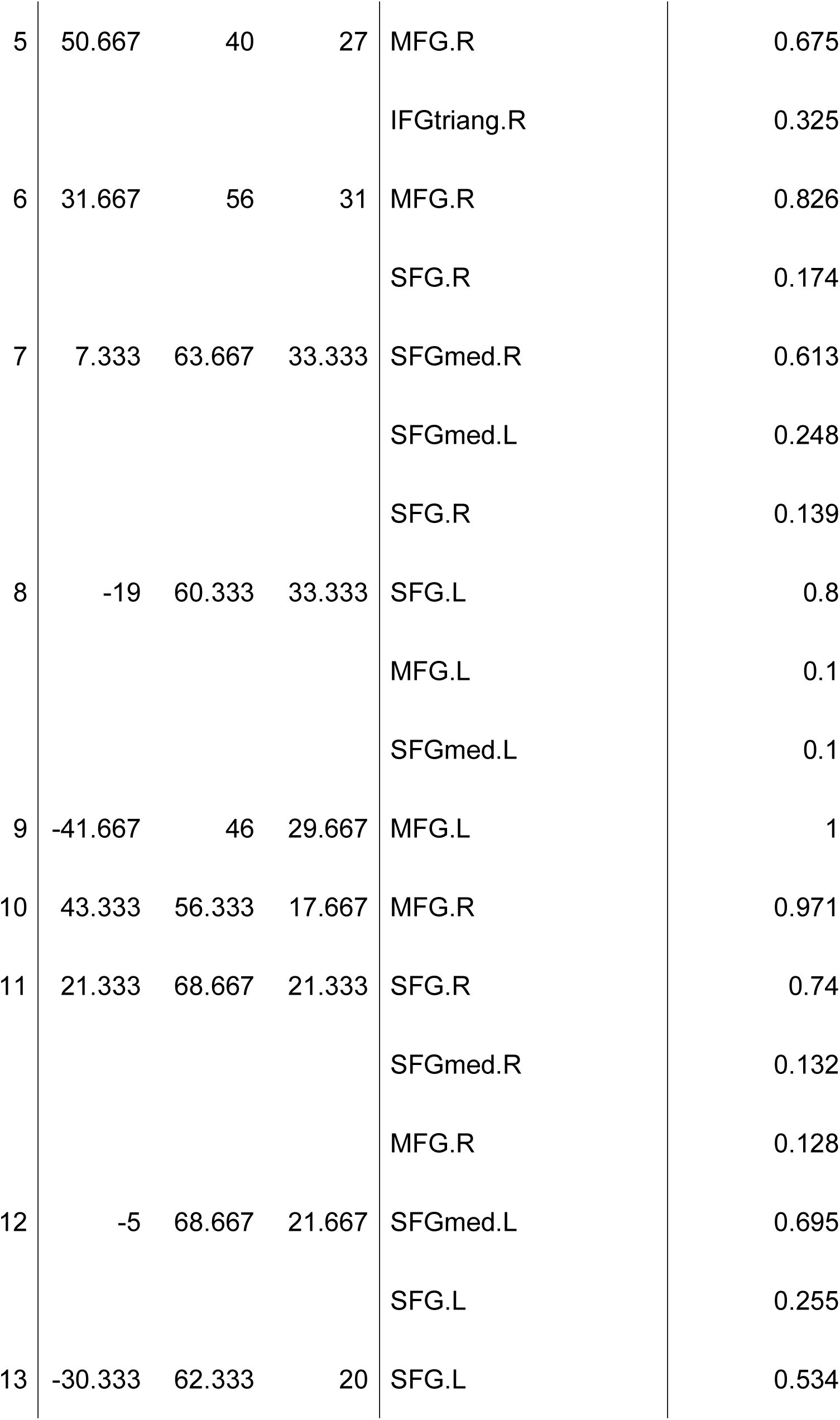
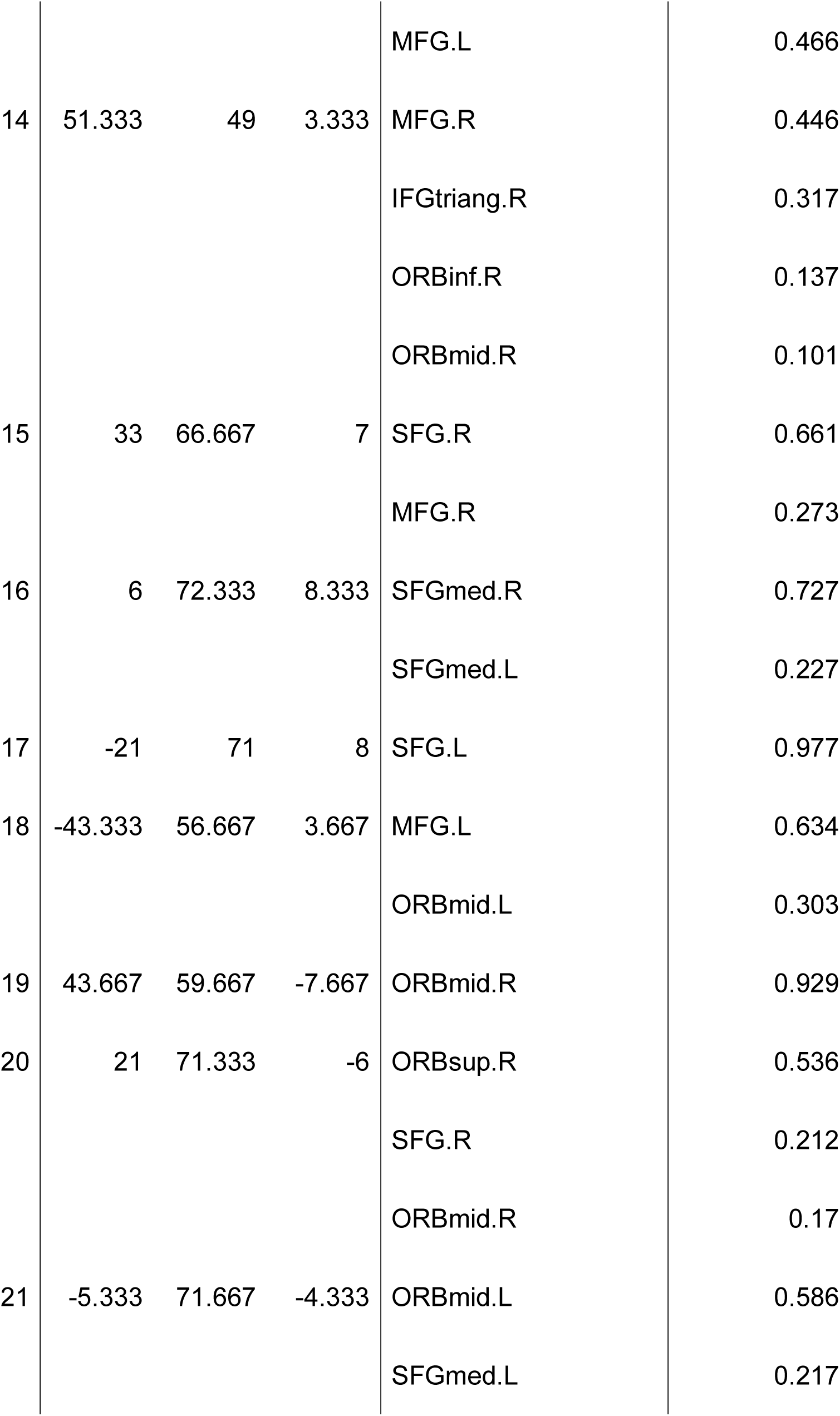
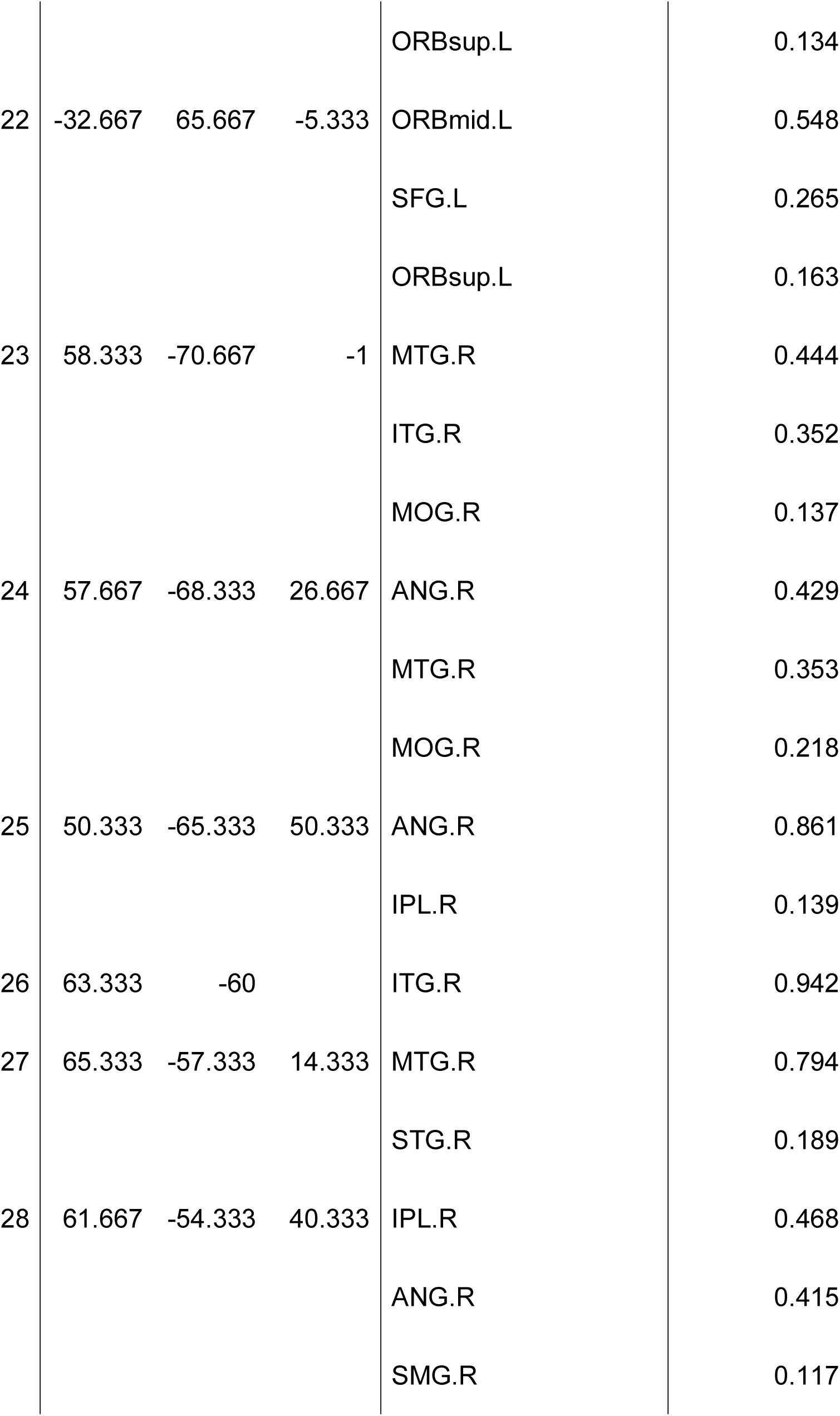
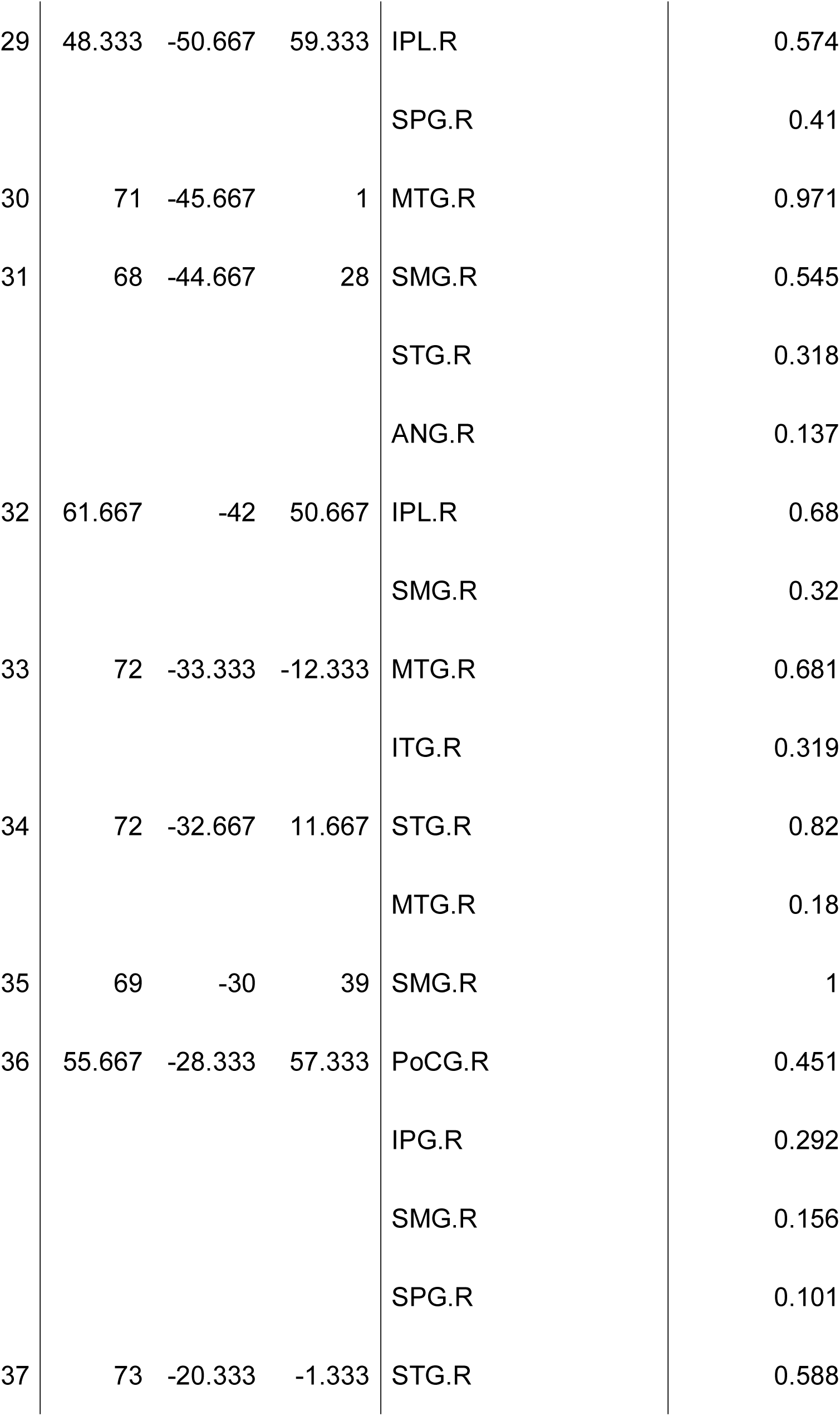
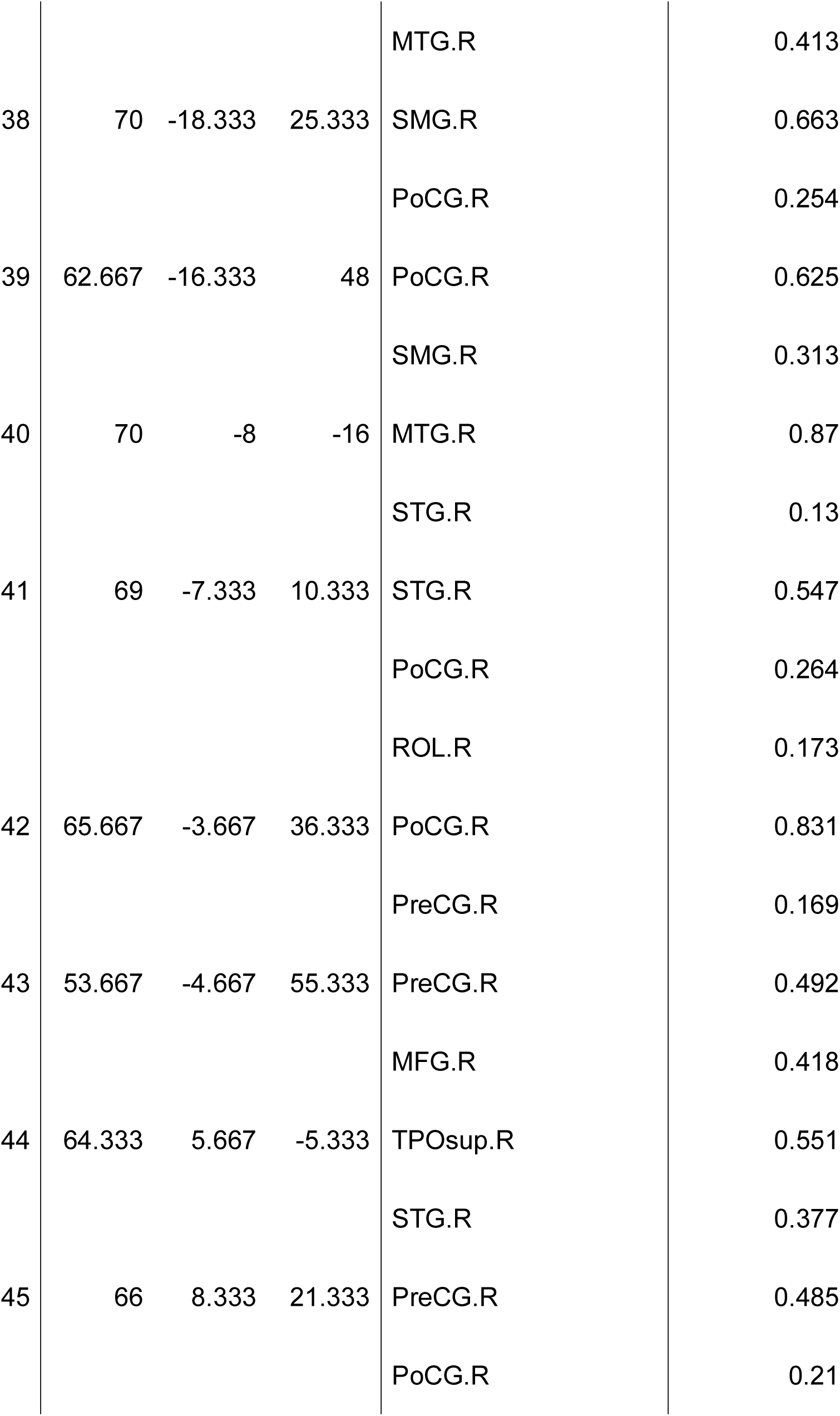
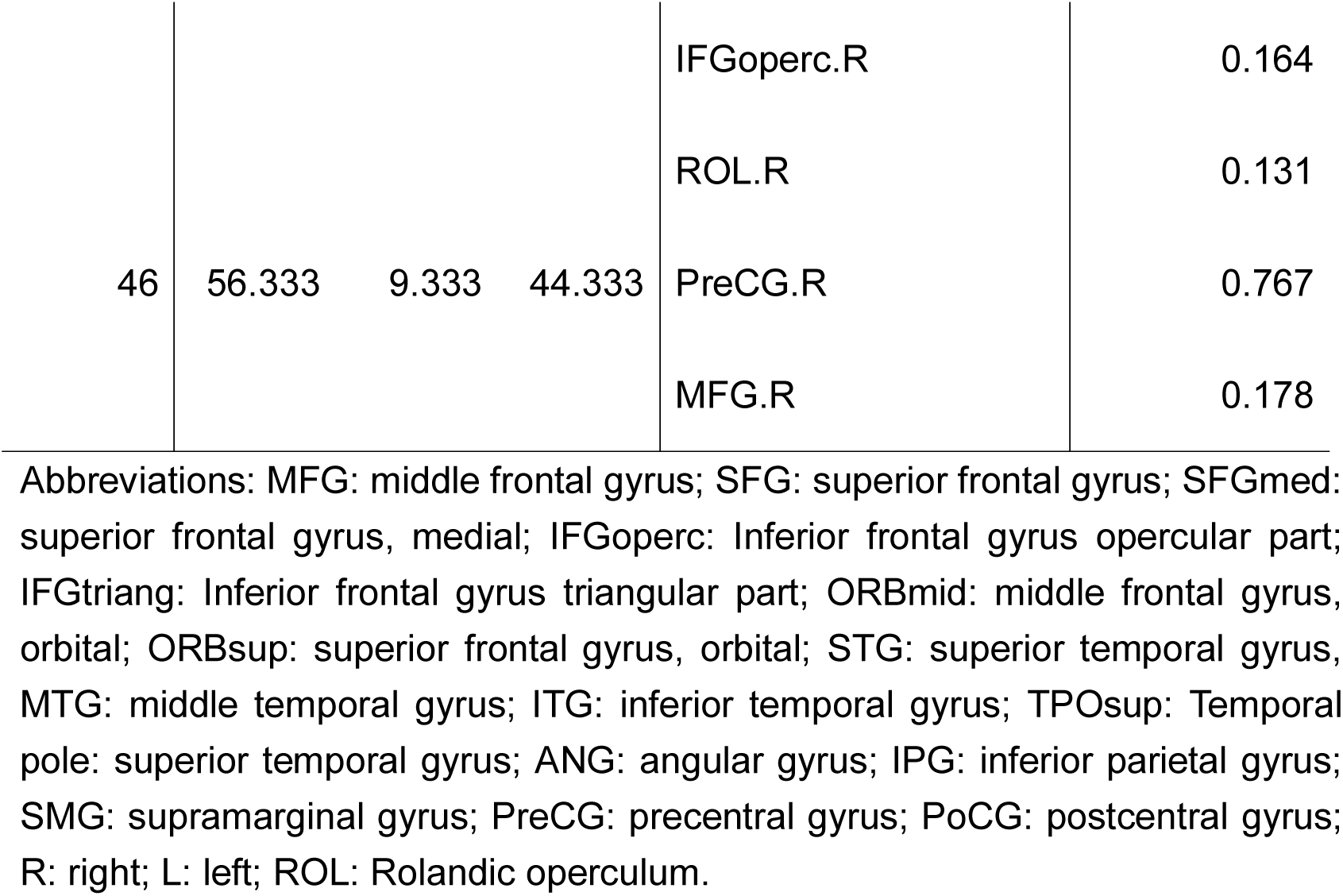
Measurement channels, group-averaged montreal neurological institute (MNI) coordinates (n = 48), anatomical regions, and region probabilities.

The task was an ultimatum game with the same design as that used by Tang et al. (2016). The difference from the original study was the use of the Japanese monetary unit in representing the reward. Before starting the experiment, each participant was randomly assigned to either the role of a Proposer or a Responder. This role was maintained until the end of the experiment. The original study added the following two stages to the traditional ultimatum game (Güth et al., 1982): (1) a stage where the Proposer tells the Responder the amount of reward (the Proposer can deceive the Responder by stating a false amount), and (2) a stage where the Proposer judges whether the Responder accepts his offer, and the Responder judges whether the Proposer has reported honestly. Adding these steps allowed us to measure how well the pairs could understand the other’s intentions.

Each pair played 54 rounds of the ultimatum game together, divided into three task blocks, each block lasting approximately 8 min. One ultimatum game consisted of six stages (Fig. 1B). In stage 1, the Proposer received one of six different rewards (¥100, ¥200, ¥500, ¥1000, ¥1500, and ¥2000). In stage 2, the Proposer tells the Responder the reward received. At this time, the Proposer was able to deceive the respondent by giving a false amount (honesty rate [HR]: the percentage of the reward reported to the Responder compared to the amount received by the Proposer). In stage 3, the Proposer decides how much of the reward is distributed to the Responder. The percentage of the amount distributed to the Responder (the offer) to the amount received by the Proposer (the actual total amount) was defined as the offer proportion (OP). In addition, the percentage of trials in which exactly 50% of the reward amount told by the Proposer to the Responder was distributed was defined as the fairness rate (FR). In stage 4, the participants evaluated each other’s actions. The Proposer judged whether the Responder would accept the offer or not, and the Responder judged whether the Proposer had honestly reported the amount received. The percentage of trials in which the pair judged each other’s behavior positively (i.e., the Proposer judged that the Responder would accept the offer and the Responder judged that the Proposer told the truth) was defined as the shared intentionality rate (SIR). In stage 5, the Responder decided whether to accept the offer. The percentage of trials in which the Responder rejected the offer was defined as the rejection rate (RR). If the Responder accepted the offer, both players received the reward as distributed by the Proposer. Otherwise, no one received rewards. In stage 6, the true winnings of each participant were shown (for example, the Proposer received ¥1000, told the Responder that the reward amount was ¥800, and distributed ¥400. In this case, the display will show “Proposer: ¥600, Responder: ¥400”). To measure emotional effects during the interaction, participants were asked to complete the Positive and Negative Affect Schedule (PANAS) scale after the experiment.

### 2.3 fNIRS data acquisition

An ETG-7100 optical topography system (Hitachi, Ltd., Tokyo, Japan) was used to record the concentration changes in oxyhemoglobin (HbO) and deoxyhemoglobin (HbR) for each pair (Fig. 1 C). The absorption of near-infrared light (wavelength: 695 ± 20 nm and 830 ± 20 nm, light intensity: 2 mW) was measured at a sampling rate of 10 Hz. Two probe sets were fitted to each subject: a 3 × 5 optode probe set (eight emitters and seven detectors, optode distance: 30 mm, 22 channels in total) and a 4 × 4 optode probe set (eight emitters and eight detectors, optode distance: 30 mm, 24 channels in total). The 3 × 5 probe set was attached to the frontal region, and the 4 × 4 probe set was attached to the right temporoparietal region according to the reference points of the International 10-20 System.

The measured data were converted into HbO and HbR concentration changes using the modified Beer–Lambert’s law. A three-dimensional digitizer (PatriotTM, Polhemus) was used to record the exact spatial coordinates of five reference points (nasion, ion, top, left ear, right ear) and 31 optical probes. To investigate the correspondence between the fNIRS channels and the brain regions, the anatomical regions (Singh et al., 2005) corresponding to each channel were estimated using the virtual registration method (Tzourio-Mazoyer et al., 2002).

### 2.4 Data analysis

#### 2.4.1 Behavioral data

The four behavioral data types (HR, OP, FR, RR) were homogeneous in variance (Bartlett’s test) and followed a normal distribution (Shapiro–Wilk test). However, the SIR was homogeneous in variance but was not normally distributed. The four behavioral data types (HR, OP, FR, RR) were analyzed using a 3 × 2 analysis of variance (ANOVA) with the time (Block1, Block2, and Block3) as a within-subject factor and the condition (FF, FB) as a between-subject factor (Bonferroni corrected). Since the SIRs did not follow a normal distribution, they were analyzed using the Friedman test instead of the 3 × 2 ANOVA. An independent sample t-test was used to test whether the total amount of rewards earned by the Proposer and Responder differed between the FF and FB conditions.

#### 2.4.2 fNIRS data: interpersonal brain synchronization

Wavelet transform coherence (WTC), defined as the cross-correlation between two time series as a function of frequency and time (Grinsted et al., 2004), has often been used to measure the IBS of fNIRS data (Tang et al., 2016; Cui et al., 2012; Jiang et al., 2012). In this study, the WTC of the HbO time series between pairs was analyzed using the MATLAB package (http://noc.ac.uk/using-science/crosswavelet-wavelet-coherence) in the frequency band between 12.8 and 51.2 s (i.e., 0.02–0.08 Hz) that was sensitive to our task. The frequency band was chosen based on the original study (Tang et al., 2016), which enabled the removal of low- and high-frequency noise from the raw HbO time series so that no additional filtering was required.

Each trial’s HbO time series from stages 2–4 (where interaction occurred between participants in the FF condition) were extracted as task data. Consequently, the task data’s duration was up to 500 s per block. The average coherence values in this band were calculated for the resting and task blocks. The relative change in the mean coherence values from the resting state to the task block was used as a measure of the change in IBS between pairs (Tang et al., 2016; Cui et al., 2012; Jiang et al., 2012). Fisher’s z-transformation was used to transform the mean coherence change for statistical tests. To find the channels where significant synchronization was observed during the task, a one-sample *t-test* (*p* < 0.05, two-tailed, false discovery rate [FDR] corrected) was performed on the coherence changes of all channels.

Next, we performed an independent samples *t*-test (*p* < 0.05, one-tailed, FDR corrected) to test whether the coherence change during the task differed between the FF and FB conditions. The *t*-values of each channel derived from these tests were used to generate the *t*-maps, which were smoothed using the spline method. Then, bivariate Pearson’s correlations between the coherence change and the SIR score were calculated to examine the association between IBS and behavior, and the calculated correlation coefficients were applied to the uncorrelated test (*p* < 0.05, FDR corrected).

Finally, we examined whether the IBS during the task was due to social interaction rather than task-derived effects. Pseudo pairs were created by randomly shuffling pairs of participants under the same conditions. This process included the following constraints: (1) the pseudo pair did not consist of the actually paired participants, and (2) the pseudo pair consisted of Proposer and Responder roles. The IBS values of the paired and pseudo paired participants were compared using permutation analysis (Nguyen et al., 2021; Bliek et al., 2015). We repeatedly generated 1000 pseudo pairs and compared their coherence changes with those of real pairs. The *p*-values were FDR-corrected for multiple comparisons (Nguyen et al., 2021; Bliek et al., 2015).

## 3. Results

### 3.1 Behavioral results

There was no significant main effect of HR, OP, and FR on both time (HR: *p* = 0.629, *F* = 0.467, η*_p_^2^* = 0.014; OP: *p* = 0.859, *F* = 0.152, η*_p_^2^* = 0.0046; FR: *p* = 0.831, *F* = 0.186, η*_p_^2^* = 0.0056) and condition (HR: *p* = 0.256, *F* = 1.313, η*_p_^2^* = 0.0195; OP: *p* = 0.436, *F* = 0.614, η*_p_^2^* = 0.0092; FR: *p* = 0.615, *F* = 0.255, η *^2^* = 0.0038). Furthermore, there was no interaction effect of condition and time on these measures. There were significant differences in the SIR and RR between the conditions (SIR, *p* = 0.003, *F* = 13.106, *η_p_^2^* = 0.191; RR: *p* = 0.047, *F* = 4.109, η*_p_*^2^ = 0.0586). However, there was neither a significant main effect of time nor a condition × time interaction effect on these behavioral measures. There was no significant difference between the conditions for the total amount of game gains (*p* = 0.43, *d* = 0.33). However, there was a significant difference in gains between the Proposer and Responder in both conditions (FF: *p* = 0.0012, *d* = 0.99; FB: *p* = 0.0003, *d* = 1.65). There was no difference in the PANAS score of each pair between the conditions (positive: *t* = -1.80, *p* = 0.077, *d* = 0.52; negative: *t* = 0.49, *p* = 0.62 *d* = 0.14).

### 3.2 fNIRS results

The results of the one-sample t-test showed that coherence did not increase significantly for all channels in the frequency range of 0.02-0.08 Hz (Fig. 3) in either condition. The independent-samples t-test showed that there was no significant difference between conditions in the coherence increases during the task. No significant correlations were found between increasing coherence and SIR, for all channels in either condition (Fig. 4).

**Fig. 3.**
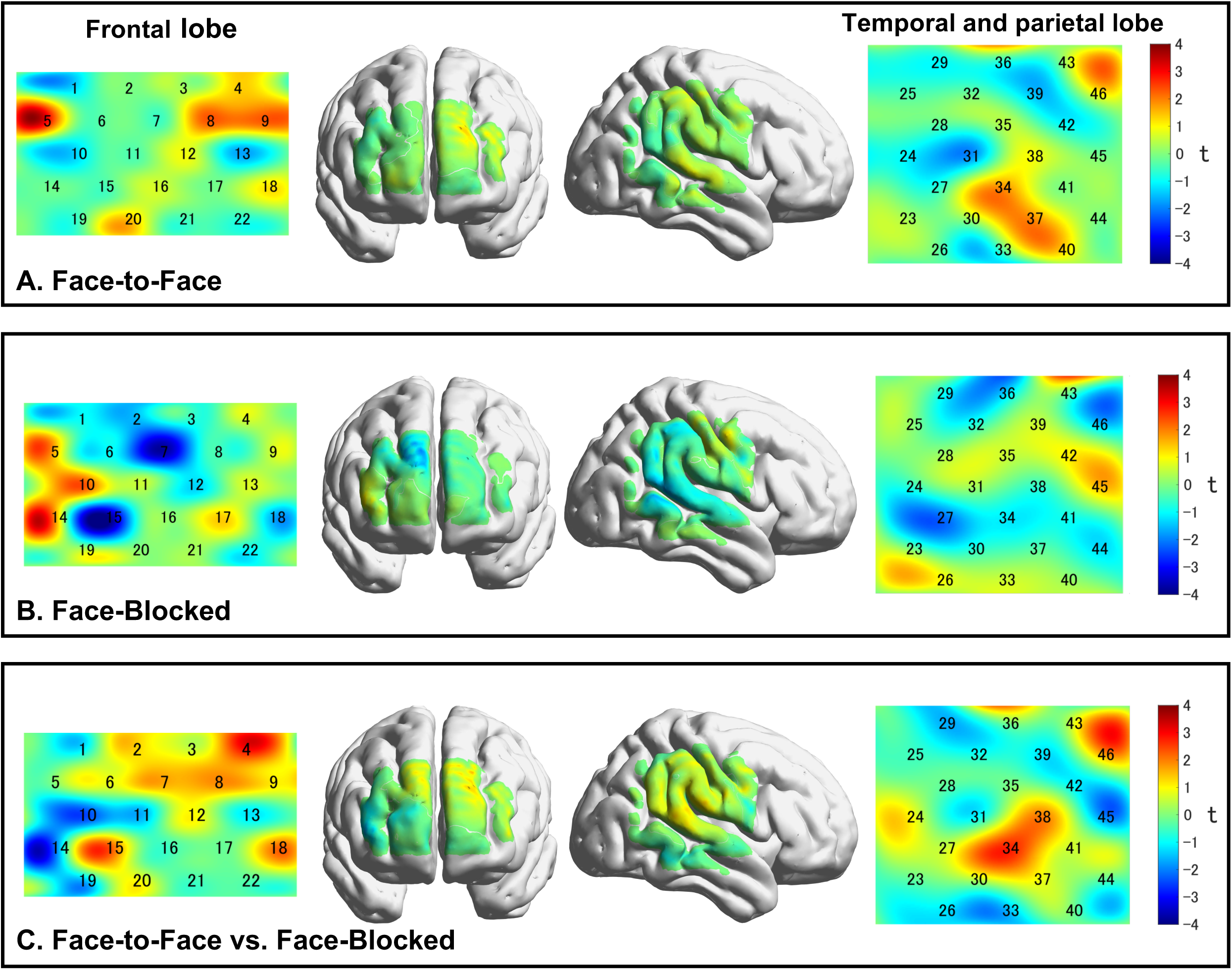
Comparison of the IBS between different conditions. (A, B) One-sample t-test for the increased coherence. In both conditions, the coherence increases in the task block from the resting state block were not significant for all channels (*p* > 0.05). (C) Independent sample t-test for the coherence increases between conditions. No conditional differences were found for all channels.

**Fig. 4.**
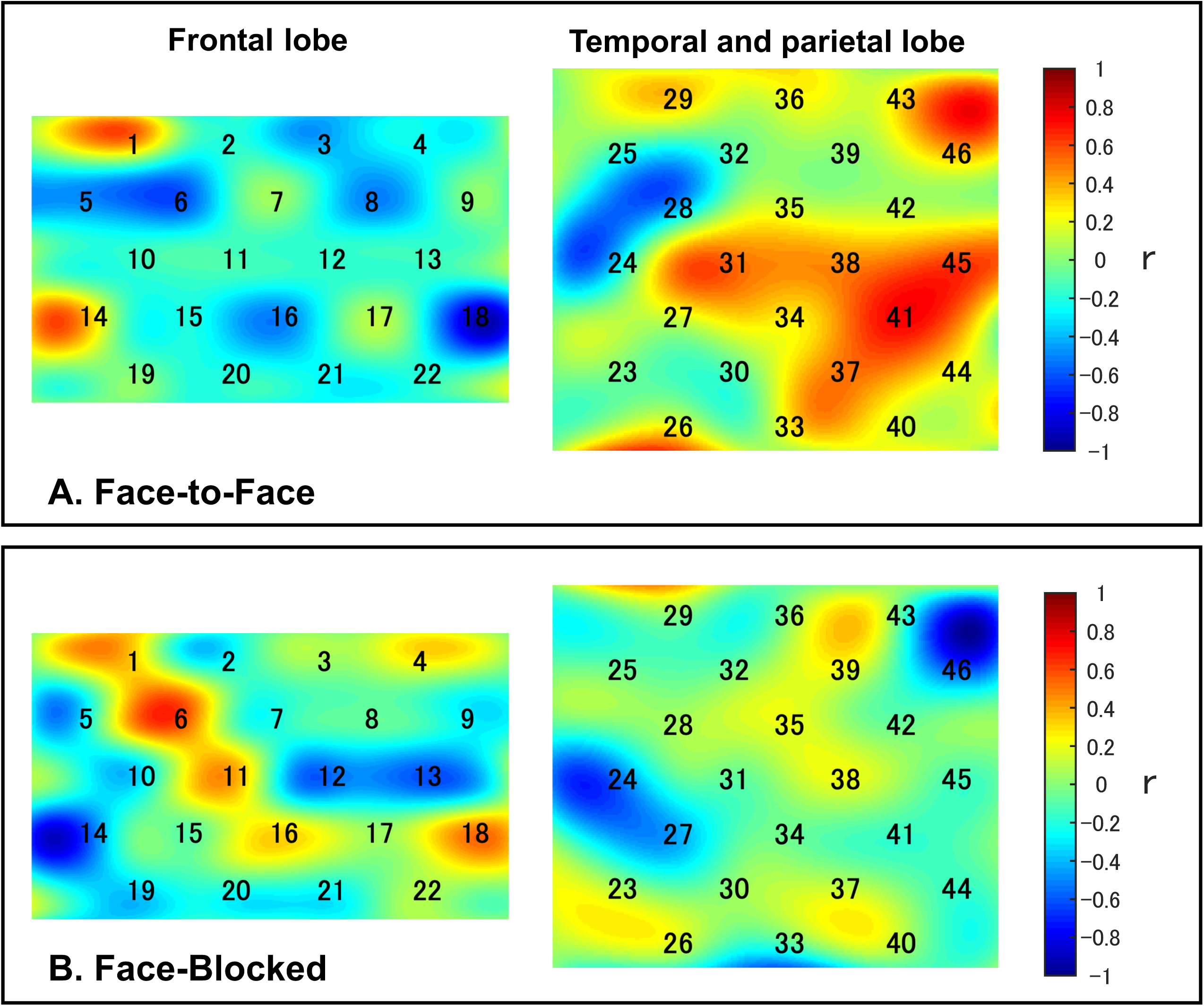
Correlations between the coherence increase and the SIR. Correlation coefficients between increased coherence and the SIR for all CHs are shown in the r-map. (A, B) There was no significant correlation between increased coherence and the SIR in either condition. Abbreviations: SIR: shared intentionality rate; CHs: channels

The results of the random pair analysis for the task block identified that the IBS of the interacting (real) pair was significantly greater than that of the pseudo pair in both conditions (Fig. 5 and Table 2). In the FF condition, there were 11 channels in which the IBS of the interacting pair was greater than that of the pseudo pair (p < 0.05, FDR-corrected): channel (CH) 4 (right superior frontal gyrus [SFG.R]), CH5 (right middle frontal gyrus [MFG.R]), CH8 (SFG.R), CH9 (MFG.R), CH12 (SFG.R), CH16 (SFG.R), CH20 (right orbital superior frontal gyrus [ORBsup.R]), CH34 (right superior temporal gyrus [STG.R]), CH36 (right postcentral gyrus [PoCG.R]), CH37 (STG.R), CH46 (right precentral gyrus [PreCG.R]). In the FB condition, two channels had significantly greater IBS values in the interacting pair than in the pseudo-pair (p < 0.05, FDR-corrected): CH10 (MFG.R) and CH14 (left orbital superior frontal gyrus [ORBsup.L]). Contrarily, the results of the random pair analysis for the resting-state block showed no significant difference in the IBS between interacting and pseudo pairs for all channels. This result was the same for both the conditions.

**Fig. 5.**
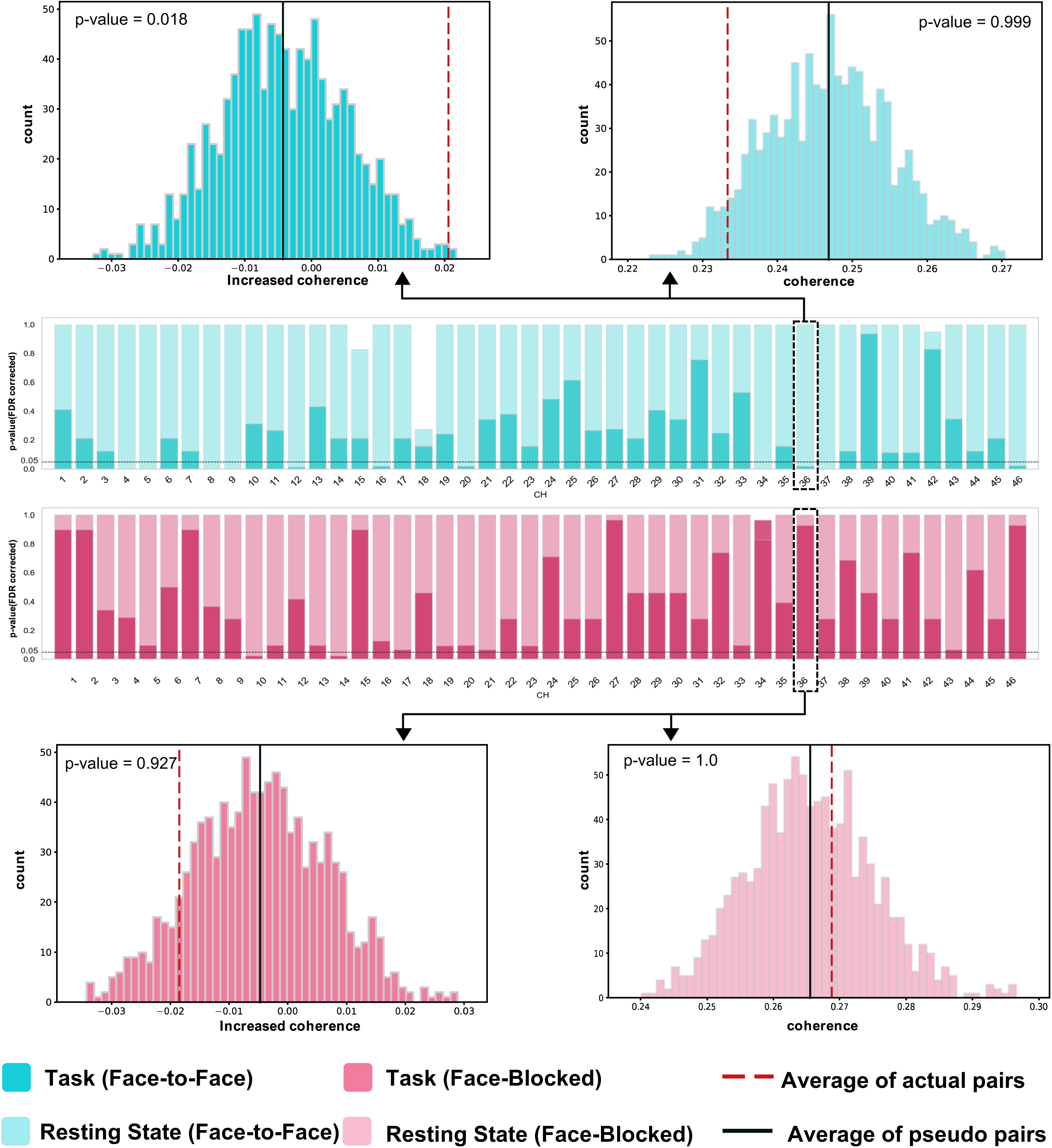
Results of the permutation test in random pair analysis (for the task and resting-state blocks). 11/46 channels indicated greater coherence in the real pair than in the pseudo pair for the task block, in the FF condition (p < 0.05, FDR corrected). In the FB condition, 2/46 channels were identified. There was no significant difference in coherence between interacting and pseudo pairs for all channels for the resting-state block. Abbreviations: FF: face-to-face; FB: face-blocked; FDR: false discovery rate

**Table 2.** The channels where the IBS of the interacting pair was significantly greater than that of the pseudo pair. Channels (CHs), conditions, and the anatomical regions associated with the CHs, p-value, and FDR-corrected p-value are listed.

| CH | Condition | Anatomical region | $p$-value | |
| --- | --- | --- | --- | --- |
| | | | $p$-value | (FDR-corrected) |
| 4 | FF | SFG.R | 0 | 0 |
| 5 | FF | MFG.R | 0 | 0 |
| 8 | FF | SFG.L | 0 | 0 |
| 9 | FF | MFG.L | 0 | 0 |
| 12 | FF | SFG.L | 0.002 | 0.013143 |
| 16 | FF | SFG.R | 0.004 | 0.0184 |
| 20 | FF | ORBsup.R | 0.003 | 0.01725 |
| 34 | FF | STG.R | 0 | 0 |
| 36 | FF | PoCG.R | 0.004 | 0.0184 |
| 37 | FF | STG.R | 0 | 0 |
| 46 | FF | PreCG.R | 0.005 | 0.020909 |
| 10 | FB | MFG.R | 0.001 | 0.023 |
| 14 | FB | ORBsup.L | 0.001 | 0.023 |

Abbreviations: MFG: left middle frontal gyrus; SFG: superior frontal gyrus; SFGmed: superior frontal gyrus, medial; ORBmid: middle frontal gyrus, orbital; ORBsup: superior frontal gyrus, orbital; STG: superior temporal gyrus, PoCG: postcentral gyrus; R: right; L: left.

## 4. Discussion

### 4.1 Behavioral results

Our results were both similar and different to those of previous studies. We succeeded in replicating the original study’s three findings. First, it was shown that the Proposer tended to distribute the reward relatively fairly, while the Responder tended to reject the offer (Fig. 2A). Previous studies have reported that the Responder accepts fair or near-fair offers (40-50%), but the RR gradually increases as the reward distribution decreases (Sanfey et al., 2003; Corradi-Dell’Acqua et al., 2013). Our result implies that the Proposer made a relatively fair offer to the Responder so that the proposal would not be rejected. Thus, it was shown that closer social distance does not affect behaviors such as fairer distribution of rewards or higher acceptance of proposals.

Second, there were no significant between-condition differences in the behavioral measures (FR, OP, and HR) associated with the Proposer (Fig. 2). In general, it is thought that Proposers distribute rewards fairly based on two motives in ultimatum games: (1) an altruistic motive due to the social norm that rewards should be distributed fairly, and (2) a strategic motive to prevent Responders from rejecting proposals (Weiland et al., 2012). A previous study has shown that disclosure of information about the Responder does not affect the Proposer’s offer and suggested that Proposers distribute rewards with strategic rather than altruistic motives (Charness et al., 2008). In the present study, the Proposer’s selfish behavior tended to be the same as in the previous study, even in acquainted pairs (Fig. 2A). Therefore, our results revealed that the FF condition or acquaintance with the partner did not affect the Proposer’s behavior regarding fairly distributing the reward.

Lastly, there were significant between-condition differences in terms of the behavioral measures of SIR and RR (Fig. 2A). Previous studies have shown that social cues play an important role in building trusting relationships with others (Chang et al., 2016; Van’t Wout et al., 2008). In the FF condition, participants could communicate nonverbally in real-time, which may have facilitated an increase in the SIR. Our results imply that the increased SIR between the pair and the Responder’s positive rating of the Proposer’s behavior made the offer more likely to be accepted.

Next, the four earlier findings of the original study were not replicated. First, in contrast to the original study, we did not find a significant main effect of time on the SIR in the FB condition (Fig. 2). In the original study, participants in the FB condition did not meet their partner directly and could only infer their partner’s personality through their behavior during the task. Therefore, since trust in the partner depends on the partner’s behavior during the task (Chang et al., 2016), it is considered that the SIR in this condition decreased over time. On the other hand, the participants in the FB condition of our study did not see their partner during the task, but they knew their identity and personality. Therefore, it can be presumed that expectations and predictions about the partner’s behavior were made based on prior information (Vavra et al., 2018; Sanfey et al., 2009). Thus it can be inferred that even in the non-face-to-face condition, the behavior during the task did not significantly affect shared intentionality. Second, the total amount of money earned by the Proposer and the Responder was not significantly different between the conditions (Fig. 2C). Third, the results of this study had a lower SIR and higher RR than those of the previous study (Fig. 2A).

These results suggest that the pairs in this study may not have cooperated as well as those in the original study. Wu et al. (2011) examined how social distance affects recipients’ evaluations of unfair behavior. They reported that in a dictatorship game, participants did not react negatively to the unfair behavior of strangers but confirmed negative reactions to the unfair behavior of their friends. This result suggests that the lesser the social distance to the other person, the more likely they are to demand fair behavior from that person. Because the social distance between the pairs in this study was closer than in the original study, it is considered that the Responder required the Proposer to distribute the reward more fairly than when the partner was a stranger. Lastly, the RR in this study was higher than that in the original study, which may indicate that the inequitable distribution accepted by the Responder in the previous study was rejected in this study. However, we could not determine whether the Responder’s motivation to reject the offer was altruistic (i.e., to correct the inequitable behavior) or selfish (i.e., to increase their reward). The frequent rejection of offers by the Responder, and the failure to understand the Responder’s intentions in rejecting the offers, possibly caused the Proposer to lose trust in the Responder. As a result, the SIR in this study may be lower than that in the original study.

### 4.2 Interpersonal brain synchronization

For IBS, the results of the original study were not replicated in our study. Here, we discuss the reasons for these differences from prior studies on IBS. In this study, there was no significant increase in coherence during the task in either the FF or FB conditions (Fig. 3A and B). Furthermore, there was no significant between-condition difference in coherence increase during the task (Fig. 3C). Previous research has shown that the lesser the social distance to the partner, the more cooperative the behavior and the greater the IBS (Pan et al., 2018).

Therefore, we expected the IBS to be greater than in previous studies because we decreased the social distance between the pairs. However, the results were contrary to our expectations, which could mean that the pairs who participated in this study did not cooperate as well as those in the previous studies. According to the behavioral analysis results, the SIR between pairs was lower, and the RR was higher in this study than in the previous study. One reason for this may be that it took a long time to come to a mutually compromised offer between the pairs because they were closer in social distance. Therefore, the IBS of acquainted pairs who did not act cooperatively may have been lower than that of stranger pairs who did cooperate.

Although the earlier IBS findings in the original study were not replicated in our acquainted pairs, these results support the idea of Gvirts et al. (2020) that IBS is influenced by with whom we interact and how. From the results of this and the original studies, the following points have been clarified. (1) If the pairs are not acquainted (i.e., the social distance between the pairs is great), FF interaction makes it easier for them to infer each other’s intentions and states of mind and cooperate. (2) Even if the pairs are acquainted (i.e., the social distance between the pairs is small), face-to-face interaction makes it easier for them to guess each other’s intention and state of mind, as in the case of the stranger pairs. However, acquaintance promotes the feeling that the partner should behave fairly, preventing them from cooperating. In summary, the results suggest that feelings toward others and the process of building shared intentionality differ depending on the social distance between members of a pair.

Furthermore, we performed random pair analysis to confirm whether the IBS during the task was specific to the interacting pair. Fig. 4 shows that the IBS of the actual pairs was significantly greater than that of the pseudo pairs. In other words, the synchronization of brain activity was not due to the execution of the same tasks between two persons but to social interaction between pairs. Therefore, the channels (brain regions) where IBS is observed are specific to the interacting pair. Previous studies have also shown that the IBS of interacting pairs is significantly larger than that of pseudo pairs (Bilek et al., 2015; Fishburn et al., 2018).

Additionally, the random pair analysis results for the resting-state block showed that the coherence of the interacting pairs was not significantly different from that of the pseudo pairs, which was the same for both conditions. This result revealed that the increase in IBS was not induced simply by being in the same space and time but only by social interaction. In addition, the brain regions where IBS in the real pairs was greater than that in the pseudo pairs were involved in social cognitive functions. In both conditions, pair-specific IBS was observed in the left and right SFG, MFG, and ORBsup. This may be because the brain functions involved in executing the ultimatum game are common to both conditions. The dorsolateral prefrontal cortex (DLPFC) is involved in working memory functions that store opponent responses during repeated strategic games (Weiland et al., 2012; Barraclough et al., 2004), and in executive control functions that control selfish and altruistic behavior (Morewedge et al., 2014; Knoch et al., 2006).

The ORBsup is involved in learning from past experiences and predicting rewards. It is presumed that participants learned about their own and others’ behavior and the associated consequences in the ultimatum game and engaged in trial and error of the optimal behavior to maximize their reward. Only in the FF condition, pair-specific IBS was observed in the STG.R, PreCG.R, and PoCG.R. This may be because social cognitive functions were more active in the FF condition. Notably, the PreCG and PoCG are involved in cognitive empathy (Seehausen et al., 2016; Takagishi et al., 2014). In ultimatum games, participants need to predict the intentions and beliefs of their opponents by taking their opponents’ point of view and control their selfish choices (e.g., unfair distribution and rejection as punishment for unfair distribution), and the IBS in the PreCG.R and PoCG.R could contribute to self-control. The STG is involved in the theory of mind, mentalizing, and the ability to understand others’ mental states (Polezzi et al., 2008; Noah et al., 2017; Carter et al., 2013). The STG, a constituent region of the TPJ, is known for integrating and processing social signals. In the FF condition, participants were able to observe their partner’s social signals (e.g., facial expressions and gestures). This may allow the participants to infer their partner’s intentions and emotions and try to understand each other’s strategies.

In summary, the participants focused their attention on their partner, memorized their actions and outcomes, and predicted the other person’s actions based on their past experiences. It is believed that they were trying to achieve the goal of maximizing rewards through these behaviors. Furthermore, FF interaction allows participants to communicate nonverbally (e.g., eye contact and gestures), making it easier for them to infer the other person’s intentions and feelings because they can receive feedback from the other person on their own actions. We identified the key regions (STG.R, PreCG.R, and PoCG.R) of the above-mentioned interaction in the FF condition using random pair analysis.

## Strengths of the current study

This study had two goals: (1) to replicate the results of Tang et al. (2016) in acquainted pairs, and (2) to demonstrate that IBS during the task is induced by social interaction. For the first objective, we found a similar trend in some behavioral data. However, the results differed from those of previous studies regarding low pairwise reliability, the high RR, and the lack of significant IBS. These results suggest that changing the nature of the interacting partners (changing the pairs from strangers to acquaintances, reducing the social distance) caused changes in the participants’ feelings toward their partners during the task and in the process of building cooperation and shared intentionality.

The results of previous studies and our behavioral data revealed that the interaction setting (FF or FB) resulted in changes in the participants’ cooperative behavior. It was confirmed that social interaction-induced IBS is affected by the nature of the interacting partners. The results of the original study by Tang et al. (2016) and the current study support the idea of Gvirts et al. (2020) that IBS is influenced by with whom we interact and how. We did not identify any significant IBS, but we did find pair-specific IBS results using random pair analysis. We succeeded in identifying such pair-specific regions: left and right SFG, MFG and ORBsup, and the right STG, PreCG, and PoCG. Furthermore, the coherence of these IBSs in the resting state did not significantly differ between the real and pseudo pairs, indicating that coherence was increased by the social interaction between the real pairs.

## Acknowledgements

We would like to thank Editage (www.editage.com) for English language editing.

## Funding

This work was supported by JSPS KAKENHI Grant Number JP20K11963.

## Notes

### Competing Interest Statement

The authors have declared no competing interest.

### Summary of Updates

F value and effect size in the behavioral result are added in Section 3.1.

## References

1. Andrews-Hanna, J. R., Smallwood, J., & Spreng, R. N. (2014). The default network and self-generated thought: component processes, dynamic control, and clinical relevance. Annals of the New York Academy of Sciences, 1316(1), 29.

2. Azhari, A., Leck, W. Q., Gabrieli, G., Bizzego, A., Rigo, P., Setoh, P., … & Esposito, G. (2019). Parenting stress undermines mother-child brain-to-brain synchrony: A hyperscanning study. Scientific reports, 9(1), 1–9.

3. Barraclough, D. J., Conroy, M. L., & Lee, D. (2004). Prefrontal cortex and decision making in a mixed-strategy game. Nature neuroscience, 7(4), 404–410.

4. Benjamini, Y., & Hochberg, Y. (1995). Controlling the false discovery rate: a practical and powerful approach to multiple testing. Journal of the Royal statistical society: series B (Methodological), 57(1), 289–300.

5. Bilek, E., Ruf, M., Schäfer, A., Akdeniz, C., Calhoun, V. D., Schmahl, C., … & Meyer-Lindenberg, A. (2015). Information flow between interacting human brains: Identification, validation, and relationship to social expertise. Proceedings of the National Academy of Sciences, 112(16), 5207–5212.

6. Boyd, R., & Richerson, P. J. (2009). Culture and the evolution of human cooperation. Philosophical Transactions of the Royal Society B: Biological Sciences, 364(1533), 3281–3288.

7. Chabin, T., Tio, G., Comte, A., Joucla, C., Gabriel, D., & Pazart, L. (2020). The relevance of a conductor competition for the study of emotional synchronization within and between groups in a natural musical setting. Frontiers in psychology, 10, 2954.

8. Chang, L. J., Doll, B. B., van’t Wout, M., Frank, M. J., & Sanfey, A. G. (2010). Seeing is believing: Trustworthiness as a dynamic belief. Cognitive psychology, 61(2), 87–105.

9. Charness, G., & Gneezy, U. (2008). What’s in a name? Anonymity and social distance in dictator and ultimatum games. Journal of Economic Behavior & Organization, 68(1), 29–35.

10. Cheng, X., Li, X., & Hu, Y. (2015). Synchronous brain activity during cooperative exchange depends on gender of partner: A fNIRS - based hyperscanning study. Human brain mapping, 36(6), 2039–2048.

11. Ciaramidaro, A., Toppi, J., Casper, C., Freitag, C. M., Siniatchkin, M., & Astolfi, L. (2018). Multiple-brain connectivity during third party punishment: an EEG hyperscanning study. Scientific Reports, 8(1), 1–13.

12. Corradi-Dell’Acqua, C., Civai, C., Rumiati, R. I., & Fink, G. R. (2013). Disentangling self-and fairness-related neural mechanisms involved in the ultimatum game: an fMRI study. Social cognitive and affective neuroscience, 8(4), 424–431.

13. Cui, X., Bryant, D. M., & Reiss, A. L. (2012). NIRS-based hyperscanning reveals increased interpersonal coherence in superior frontal cortex during cooperation. Neuroimage, 59(3), 2430–2437.

14. Delgado, M. R., Frank, R. H., & Phelps, E. A. (2005). Perceptions of moral character modulate the neural systems of reward during the trust game. Nature neuroscience, 8(11), 1611–1618.

15. Declerck, C. H., & Boone, C. (2018). The neuroeconomics of cooperation. Nature Human Behaviour, 2(7), 438–440.

16. Dumas, G. (2011). Towards a two-body neuroscience. Communicative & integrative biology, 4(3), 349–352.

17. Fishburn, F. A., Murty, V. P., Hlutkowsky, C. O., MacGillivray, C. E., Bemis, L. M., Murphy, M. E., … & Perlman, S. B. (2018). Putting our heads together: interpersonal neural synchronization as a biological mechanism for shared intentionality. Social cognitive and affective neuroscience, 13(8), 841–849.

18. Giorgetta, C., Grecucci, A., Graffeo, M., Bonini, N., Ferrario, R., & Sanfey, A. G. (2021). Expect the Worst! Expectations and Social Interactive Decision Making. Brain sciences, 11(5), 572.

19. Grinsted, A., Moore, J. C., & Jevrejeva, S. (2004). Application of the cross wavelet transform and wavelet coherence to geophysical time series. Nonlinear processes in geophysics, 11(5/6), 561–566.

20. Güth, Werner, Rolf Schmittberger, and Bernd Schwarze. “An experimental analysis of ultimatum bargaining.” Journal of economic behavior & organization 3.4 (1982): 367–388.

21. Gvirts, H. Z., & Perlmutter, R. (2020). What guides us to neurally and behaviorally align with anyone specific? A neurobiological model based on fNIRS hyperscanning studies. The Neuroscientist, 26(2), 108–116.

22. Hirsch, J., Zhang, X., Noah, J. A., & Ono, Y. (2017). Frontal temporal and parietal systems synchronize within and across brains during live eye-to-eye contact. NeuroImage, 157, 314–330.

23. Jiang, J., Dai, B., Peng, D., Zhu, C., Liu, L., & Lu, C. (2012). Neural synchronization during face-to-face communication. Journal of Neuroscience, 32(45), 16064–16069.

24. Knoch, D., Pascual-Leone, A., Meyer, K., Treyer, V., & Fehr, E. (2006). Diminishing reciprocal fairness by disrupting the right prefrontal cortex. science, 314(5800), 829–832.

25. Koike, T., Sumiya, M., Nakagawa, E., Okazaki, S., & Sadato, N. (2019). What makes eye contact special? Neural substrates of on-line mutual eye-gaze: a hyperscanning fMRI study. Eneuro, 6(1).

26. Koike, T., Tanabe, H. C., Adachi-Abe, S., Okazaki, S., Nakagawa, E., Sasaki, A. T., … & Sadato, N. (2019). Role of the right anterior insular cortex in joint attention-related identification with a partner. Social cognitive and affective neuroscience, 14(10), 1131–1145.

27. Liu, N., Mok, C., Witt, E. E., Pradhan, A. H., Chen, J. E., & Reiss, A. L. (2016). NIRS-based hyperscanning reveals inter-brain neural synchronization during cooperative Jenga game with face-to-face communication. Frontiers in human neuroscience, 10, 82.

28. Lu, C. M., Zhang, Y. J., Biswal, B. B., Zang, Y. F., Peng, D. L., & Zhu, C. Z. (2010). Use of fNIRS to assess resting state functional connectivity. Journal of neuroscience methods, 186(2), 242–249.

29. Lu, K., Xue, H., Nozawa, T., & Hao, N. (2019). Cooperation makes a group be more creative. Cerebral Cortex, 29(8), 3457–3470.

30. Mandel, D. R. (2006). Economic transactions among friends: Asymmetric generosity but not agreement in buyers’ and sellers’ offers. Journal of Conflict Resolution, 50(4), 584–606.

31. Morewedge, C. K., Krishnamurti, T., & Ariely, D. (2014). Focused on fairness: Alcohol intoxication increases the costly rejection of inequitable rewards. Journal of Experimental Social Psychology, 50, 15–20.

32. Müller, V., & Lindenberger, U. (2019). Dynamic orchestration of brains and instruments during free guitar improvisation. Frontiers in integrative neuroscience, 13, 50.

33. Nguyen, T., Schleihauf, H., Kungl, M., Kayhan, E., Hoehl, S., & Vrtička, P. (2021). Interpersonal Neural Synchrony During Father–Child Problem Solving: An fNIRS Hyperscanning Study. Child Development.

34. Noah, J. A., Dravida, S., Zhang, X., Yahil, S., & Hirsch, J. (2017). Neural correlates of conflict between gestures and words: a domain-specific role for a temporal-parietal complex. PLoS One, 12(3), e0173525.

35. Osaka, N., Minamoto, T., Yaoi, K., Azuma, M., Shimada, Y. M., & Osaka, M. (2015). How two brains make one synchronized mind in the inferior frontal cortex: fNIRS-based hyperscanning during cooperative singing. Frontiers in psychology, 6, 1811.

36. Overgaauw, S., Güroğlu, B., & Crone, E. A. (2012). Fairness considerations when I know more than you do: developmental comparisons. Frontiers in psychology, 3, 424.

37. Pan, Y., Cheng, X., Zhang, Z., Li, X., & Hu, Y. (2017). Cooperation in lovers: an f NIRS-based hyperscanning study. Human brain mapping, 38(2), 831–841.

38. Polezzi, D., Daum, I., Rubaltelli, E., Lotto, L., Civai, C., Sartori, G., & Rumiati, R. (2008). Mentalizing in economic decision-making. Behavioural brain research, 190(2), 218–223.

39. Rolls, E. T., Huang, C. C., Lin, C. P., Feng, J., & Joliot, M. (2020). Automated anatomical labelling atlas 3. Neuroimage, 206, 116189.

40. Saito, D. N., Tanabe, H. C., Izuma, K., Hayashi, M. J., Morito, Y., Komeda, H., … & Sadato, N. (2010). “Stay tuned”: inter-individual neural synchronization during mutual gaze and joint attention. Frontiers in integrative neuroscience, 4, 127.

41. Salazar, M., Shaw, D. J., Gajdoš, M., Mareček, R., Czekóová, K., Mikl, M., & Brázdil, M. (2021). You took the words right out of my mouth: Dual-fMRI reveals intra-and inter-personal neural processes supporting verbal interaction. NeuroImage, 228, 117697.

42. Sanfey, A. G. (2009). Expectations and social decision-making: biasing effects of prior knowledge on Ultimatum responses. Mind & Society, 8(1), 93–107.

43. Sanfey, A. G., Rilling, J. K., Aronson, J. A., Nystrom, L. E., & Cohen, J. D. (2003). The neural basis of economic decision-making in the ultimatum game. Science, 300(5626), 1755–1758.

44. Seehausen, M., Kazzer, P., Bajbouj, M., Heekeren, H. R., Jacobs, A. M., Klann-Delius, G., … & Prehn, K. (2016). Effects of empathic social responses on the emotions of the recipient. Brain and Cognition, 103, 50–61.

45. Singh, A. K., Okamoto, M., Dan, H., Jurcak, V., & Dan, I. (2005). Spatial registration of multichannel multi-subject fNIRS data to MNI space without MRI. Neuroimage, 27(4), 842–851.

46. Stallen, M., & Sanfey, A. G. (2013). The cooperative brain. The Neuroscientist, 19(3), 292–303.

47. Takagishi, H., Koizumi, M., Fujii, T., Schug, J., Kameshima, S., & Yamagishi, T. (2014). The role of cognitive and emotional perspective taking in economic decision making in the ultimatum game. PloS one, 9(9), e108462.

48. Tang, H., Mai, X., Wang, S., Zhu, C., Krueger, F., & Liu, C. (2016). Interpersonal brain synchronization in the right temporo-parietal junction during face-to-face economic exchange. Social cognitive and affective neuroscience, 11(1), 23–32.

49. Tang, H., Zhang, S., Jin, T., Wu, H., Su, S., & Liu, C. (2019). Brain activation and adaptation of deception processing during dyadic face-to-face interaction. Cortex, 120, 326–339.

50. Tzourio-Mazoyer, N., Landeau, B., Papathanassiou, D., Crivello, F., Etard, O., Delcroix, N., … & Joliot, M. (2002). Automated anatomical labeling of activations in SPM using a macroscopic anatomical parcellation of the MNI MRI single-subject brain. Neuroimage, 15(1), 273–289.

51. Van’t Wout, M., & Sanfey, A. G. (2008). Friend or foe: The effect of implicit trustworthiness judgments in social decision-making. Cognition, 108(3), 796–803.

52. Vavra, P., Chang, L. J., & Sanfey, A. G. (2018). Expectations in the Ultimatum Game: distinct effects of mean and variance of expected offers. Frontiers in psychology, 9, 992.

53. Weiland, S., Hewig, J., Hecht, H., Mussel, P., & Miltner, W. H. (2012). Neural correlates of fair behavior in interpersonal bargaining. Social Neuroscience, 7(5), 537–551.

54. Weiß, M., Paelecke, M., & Hewig, J. (2021). In Your Face (t)—Personality Traits Interact With Prototypical Personality Faces in Economic Decision Making. Frontiers in psychology, 12.

55. Wu, Y., Leliveld, M. C., & Zhou, X. (2011). Social distance modulates recipient’s fairness consideration in the dictator game: An ERP study. Biological psychology, 88(2-3), 253–262.

56. Zhang, M., Jia, H., & Zheng, M. (2020). Interbrain synchrony in the expectation of cooperation behavior: a hyperscanning study using functional near-infrared spectroscopy. Frontiers in psychology, 11.

